# Developmental and Early-Life Stress-Induced Effects on 5-HT_3_R-Expressing Interneurons within Auditory Cortex

**DOI:** 10.1101/2025.09.08.675013

**Authors:** James T. Moore, Matthew J. Sunthimer, Ethan White, Jeffrey G. Mellott, Merri J. Rosen

## Abstract

Early life stress (ELS) is a well-known predictor of neuropsychiatric disease and contributes to the development of sensory processing deficits that persist throughout life. Organisms are particularly susceptible to the deleterious effects of stress during critical periods, when neuroplasticity is heightened, and initial representations of the sensory environment are mapped to cortex. When ELS is induced during the auditory cortical (ACx) critical period, it impairs both neural and behavioral responses to a variety of auditory stimuli that rely on temporal processing. Mechanisms by which ELS may alter critical period plasticity are of particular interest in understanding ELS-related pathology, including the 5-HT3R interneuron system, which has been implicated in regulating neural activity during critical periods. Here we examined two principal subpopulations of interneurons in primary ACx: VIP and NDNF cells, which account for a majority of cortical neurons expressing 5-HT3R. The expression of the Htr3a gene during normal development and under ELS conditions was quantified using multiplex fluorescent in situ hybridization. We show that densities of cells expressing NDNF and VIP decrease following ear opening and across the ACx critical period, and that ELS results in the maintenance of elevated cell densities compared to age-matched controls. Further, Htr3a in VIP neurons is developmentally upregulated, and its expression is further increased by ELS beyond normal physiologic levels. Stress-induced shifts in these serotonergic interneurons may contribute to deficits that arise in auditory cortical and perceptual responses via effects on local cortical circuitry.

## INTRODUCTION

Early life stress (ELS) contributes to neuropsychiatric disease in both humans and animal models (Kessler et al., 2010; Chen and Baram, 2016; Yajima et al., 2018). Emotional and cognitive regions of the brain are particularly susceptible to insults from stress during ‘critical periods’: distinctive time windows of heightened plasticity where susceptibility to early adverse experiences is maximal (Dunn et al., 2019; Gabard-Durnam and McLaughlin, 2019; Malave et al., 2022). Recent investigations of ELS have broadened beyond emotional and cognitive regions to include sensory cortices, indicating that ELS can directly disrupt sensory processing (Takatsuru and Koibuchi, 2015; Liu et al., 2020; Ye et al., 2023; Poplawski et al., 2024; An et al., 2025; Mazi et al., 2025; Rosen and Huyck, 2025). When ELS is induced during the auditory cortical (ACx) critical period, it impairs both neural and behavioral responses to a variety of auditory stimuli that rely on temporal processing, such as accurate detection of brief gaps in sound, frequency modulations, and sound locations (Ye et al., 2023; An et al., 2025; Mazi et al., 2025). Mechanistically, these deficits are accompanied by reductions in several cortical elements known to be involved in critical period plasticity: inhibitory parvalbumin neurons (PV), perineuronal nets (PNNs), and brain-derived neurotrophic factor (BDNF) (Bath et al., 2013; Takesian and Hensch, 2013; Castillo-Gomez et al., 2017; Murthy et al., 2019; Guadagno et al., 2021; Sunthimer et al., 2023; An et al., 2025; Mazi et al., 2025). An important player that has received less attention is serotonin (5-HT).

Serotonin is linked to stress, developmental plasticity, and the auditory system. Serotonergic neurons project to ACx and other auditory regions, releasing serotonin based on behavioral contexts including stress (Campbell et al., 1987; DeFelipe et al., 1991; Clement et al., 1998; Kranz et al., 2010; Hurley and Hall, 2011; Koyama et al., 2017). Stress can induce widespread effects on the serotonergic system, altering levels of 5-HT and 5-HT receptors (5-HTRs) throughout the brain (Veenema, 2009; Bravo et al., 2014; Lages et al., 2021). Within the auditory system, such serotonergic shifts can affect neural excitability (Lee et al., 2010; Rao et al., 2010), sound-evoked response properties (Hurley and Pollak, 2005; Hall et al., 2010; Cheng et al., 2023b; Horrocks et al., 2025), and behavioral responses to sound (Adler et al., 2005; Tsitsipa et al., 2022). Serotonergic effects are pronounced during early postnatal development, where manipulations that increase 5-HT levels cause later behavioral hypersensitivity to sound and disruption of ACx sound-evoked response properties (Kahne et al., 2002; Simpson et al., 2011; Pan et al., 2021). Such early susceptibility suggests a role for 5-HT during critical period plasticity. Indeed, blocking 5-HT activity reduces plasticity in prefrontal, somatosensory, and visual cortices specifically during critical periods involving those regions (Cases et al., 1996; Wang et al., 1997; Rebello et al., 2014). Even in humans, increasing 5-HT levels perinatally via serotonin reuptake inhibitors accelerates the maturation of speech perception in infants (Weikum et al., 2012).

Within ACx, superficial interneurons that express 5-HT3 receptors (5-HT3R) may have a role in regulating critical period plasticity. These cells integrate bottom-up auditory information with top-down neuromodulatory input, responding to sound with response properties that are modulated by behavioral state (Sweeney et al., 2025; Vattino et al., 2025). Optogenetic inactivation of 5-HT3R cells during the ACx critical period eliminates tonotopic map plasticity in A1, demonstrating their involvement in experience-dependent developmental plasticity (Takesian et al., 2018). Converging lines of research also suggest that ELS affects 5-HT3R gene expression. In rats, 5-HT3aR mRNA levels were increased by ELS in prefrontal cortex and hippocampus (Zolfaghari et al., 2021), and in psychiatric patients, childhood maltreatment was associated with epigenetic changes in the gene encoding 5-HT3R (Perroud et al., 2016).

The extent to which 5-HT3R expression contributes to critical period plasticity or to altered circuit dynamics resulting from ELS is currently limited by a lack of detailed information concerning the native expression profiles of 5-HT3R within the two major subpopulations of 5-HT3R cells: vasoactive intestinal peptide (VIP) cells that are densely populated in superficial cortical layers, and neuron derived neurotrophic factor (NDNF) cells that are mostly confined to L1 (Kullander and Topolnik, 2021; Hartung et al., 2024). Both groups innervate a combination of other interneurons and dendrites of pyramidal cells to form complex disinhibitory circuits that are involved in gating cortical responses and guiding neuroplasticity (Donato et al., 2013; Takesian et al., 2018; Hartung et al., 2024) (Figure 1A). Here, we characterized the cell type-specific expression profiles of the Htr3a gene in these cells within A1 during normal development and under the influence of ELS. This revealed that Htr3a is differentially expressed between NDNF and VIP cell types in a layer-specific manner. Further, ELS affects both cell density and Htr3a expression levels, potentially causing long-term alterations to cortical function and neuroplasticity.

**Figure 1.**
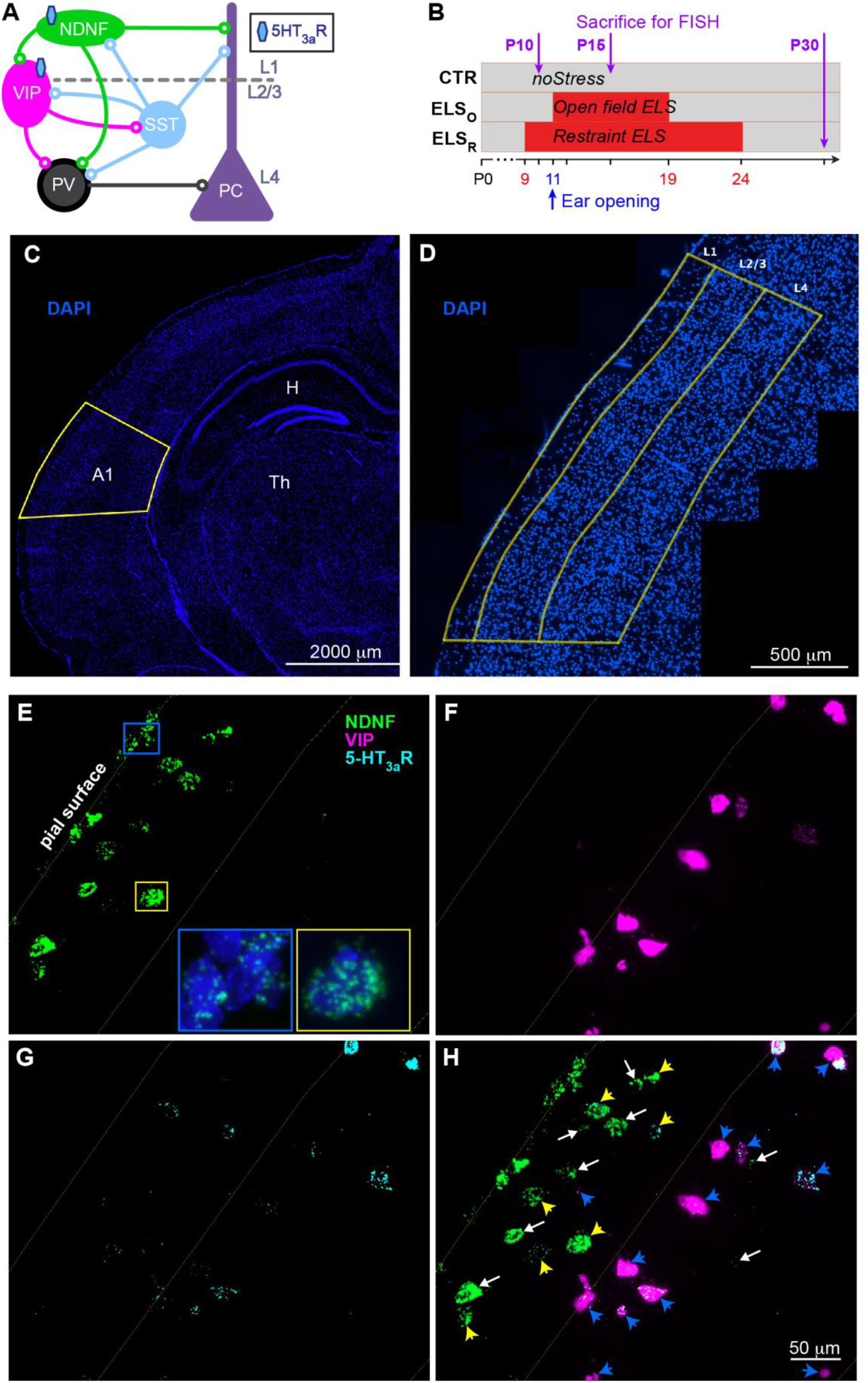
Summary of methods. (A) Simplified schematic of NDNF^+^ and VIP^+^ connectivity in ACx showing relationships to parvalbumin (PV) and somatostatin (SST) interneurons and pyramidal cells (PC). (B) Timeline of stress induction with endpoints denoting tissue collection. (C) A representative coronal section imaged at 5x in the DAPI channel identifying A1. (D) Tiled image of A1 sampling area taken at 40x, delineating L1, 2/3 and 4. (E-G) Separate fluorescent channels showing target probes for *NDNF, VIP* and *Htr3a*. Insets show *NDNF* localized to non-neuronal cells in the pial surface [blue outline] and an NDNF+ neuron [yellow outline]. (H) Composite image showing results of multichannel FISH labeling. All counted cells are indicated by arrows or arrowheads. Arrowheads depict cells with *Htr3a* puncta (yellow for NDNF cells, blue for VIP cells). White arrows depict cells with no *Htr3a* puncta (NDNF cells only).

## MATERIALS AND METHODS

### Subjects

All experiments followed procedures approved by the Northeast Ohio Medical University Institutional Animal Care and Use Committee. Mixed-sex groups of Mongolian gerbils (*Meriones unguiculatus)* were used, as they are an established animal model for the study of auditory processing. Gerbils from multiple litters were housed with littermates in a 12/12 h light/dark cycle and fed ad libitum. Control and ELS animals were collected from separate litters to avoid stressful effects on control animals or parents induced by removing siblings for ELS treatment.

### Experimental Design

Development of NDNF^+^ and VIP^+^ cell populations was examined by sampling brain sections containing A1 from five animal groups (Figure 1B). We performed mRNA *in situ* hybridization in three channels simultaneously, using custom probes targeting *NDNF, VIP*, and *Htr3a*. Control animals were sacrificed at postnatal day (P)10 and P15 to characterize a timeline of normal development in A1 during the ACx critical period, which occurs approximately from ear canal opening ∼P11 through P23 (Mowery et al., 2015). A third control group was sacrificed at P30 following closure of the critical period. Two ELS groups, “Restraint” ELS (ELS_R_) and “Open field” ELS (ELS_O_) were subjected to separate forms of stress induction during the ACx critical period as described below, and sacrificed at P30 to elucidate the residual effects of ELS on A1 development. All groups contained 6 subjects: P15 controls (CTR15’s) contained 4 males and 2 females; P30 controls (CTR30’s) contained 2 males and 4 females; all other groups were balanced equally by sex. For each group, three measures were quantified: 1) Cell population density within each layer of A1 for *NDNF* and *VIP*-labeled cells, 2) number of *Htr3a* mRNA within each NDNF and VIP cell, and 3) number of NDNF puncta within each NDNF cell.

### Stress Induction

ELS_R_ animals were subjected to intermittent sessions of maternal separation and restraint across a 16-day time window encompassing the critical period for maturation of ACx intrinsic and synaptic properties (Mowery et al., 2015). This corresponded to P9-P24 and consisted of ten 2-h restraint sessions occurring in the morning, afternoon, or evening. Time of day was varied randomly to reduce predictability in order to increase stress (Baram et al., 2012). During each session, animal cages were transported from the animal facility to a testing room, where the pups were removed and placed into 50-ml centrifuge tubes isolated from their parents and littermates. The tubes were perforated to allow easy breathing and placed either vertically or horizontally inside sound attenuated, anechoic booths with lights on. Pups were returned to their home cage and then to the animal facility immediately following each session.

ELS_O_ animals were subjected to intermittent sessions of maternal separation and isolation in open spaces during a 9-day time window corresponding to P11-P19, consisting of six 2-h open isolation sessions occurring in the morning, afternoon, or evening. The ELS_O_ group received fewer isolation sessions over a compacted developmental time window compared to the ELS_R_ group. These parameters were chosen to span the developmental time window of maximal plasticity while avoiding malnutrition and neglect, because using the timing of ELS_R_ isolation in the ELS_O_ paradigm resulted in reduced body weight (data not shown). Pups were separated from their home cage and transported to a testing room where they were placed individually into 10.5” x 12.5” clear acrylic boxes with no bedding or environmental stimuli. The boxes were then placed into sound attenuated, anechoic booths with lights off. Pups were returned to the animal facility and back to their original home cage immediately following each session.

### RNA fluorescent in situ hybridization (FISH)

Gerbils at P10, P15, or P30 were anaesthetized with isoflurane and decapitated. Brains were rapidly removed, embedded in Optimal Cutting Temperature medium (OCT, Scigen) and flash frozen. 14μm coronal sections containing A1 were then collected on a cryostat microtome (Leica CM 1950) and adhered to SuperFrost Plus slides (Fisher Scientific). Sections were chosen based on anatomical landmarks corresponding to Plates 30-32 from the brain atlas of the Mongolian gerbil (Radtke-Schuller et al., 2016). These landmarks included the presence of lateral ventricles lateral to the thalamus, and hippocampus expanding dorsoventrally. Sections were chosen to be anterior to the superior collicus and taken from the widest portion of the brain measured medio-laterally. Slides were stored under airtight conditions at - 80°C until use. *In situ* hybridization was performed using the RNAscope Multiplex Fluorescent Reagent Kit v2 (ACDBio) according to manufacturer instructions. Briefly, slides were post-fixed for 1 hour in buffered formalin, dehydrated in a series of ethanol baths (50%,70%, 100%, 100%), then pre-processed with hydrogen peroxide (10 minutes) followed by protease IV (30 minutes). RNAscope probes were hybridized at 40°C for 2 hours, then conjugated to a series of signal-amplifying molecular scaffolds, followed by fluorescent labeling using channel-specific horseradish peroxidases and TSA vivid fluorophores in 1:1500 dilution (Tocris). The probes used were as follows: Mun-Htr3a-C1(Cat. #1299361-C1), Mun-Ndnf-C2(Cat. #1299341-C2), and Mun-Vip-C3(Cat. #1299351-C3). All slides were counterstained with DAPI for 30s prior to coverslipping with Prolong Gold antifade mountant (Invitrogen).

### Imaging and Cell Quantification

Data for VIP^+^ cells were collected from a single tissue section from each subject. In addition to the primary tissue section, NDNF^+^ data were also collected from a second section, when available, to ensure sufficient sampling due to their lower abundance. A1 was identified using thalamic and hippocampal anatomical landmarks guided by the gerbil brain atlas (Figure 1C; Radtke-Schuller et al., 2016). The dorsal extent of A1 lies near the level of the dentate gyrus (DG); the region extends ventrally to a point just inferior of the lateral pole of cortex in Mongolian gerbils, ∼1-2mm superior to the rhinal fissure. We sampled a conservative area extending 2mm along the D-V axis from the level of DG, extending ventrally to just superior of the lateral pole. Both cell populations were sampled throughout neocortical layers 1-4 (L1, L2/3 and L4), where the majority of cortical 5HT3R interneurons are concentrated and have been best characterized in the existing literature (Lee et al., 2010; Kullander and Topolnik, 2021). Layers were identified visually based on packing densities illuminated by DAPI, including the koniocortical patterning of L4 seen in primary sensory areas (Radtke-Schuller et al., 2016). Tiled images of A1 containing L1-L4 were taken at 40x in four channels to generate A1 “maps” which were used to identify all cells that expressed *NDNF* or *VIP* mRNA within the region of interest (Figure 1D). We only counted cells that showed at least 2 puncta tagged with *NDNF* (NDNF^+^) or *VIP* (VIP^+^) label that were localized to an intact DAPI-stained nucleus. Cells with splintered nuclei were likely either damaged by sectioning or programmed cell death and were not counted. Because NDNF is also expressed in epithelial structures such as blood vessels, presumably in non-neuronal cells (Ohashi et al., 2014), NDNF^+^ cells that were touching the pial surface in L1 or embedded within apparent blood vessels were excluded from analysis (Figure 1E, *blue inset*). The A1 maps were subsequently used to draw layer boundaries and record area measurements using the open-source software Fiji (Schindelin et al., 2012) to assess layer distribution. Within each layer, and cell density. All imaging was performed on a Zeiss Axio Imager M2 fluorescent microscope using the accompanying ZEN Blue software. Photos were taken using a Hamamatsu model C13440 digital camera.

### Htr3a Quantification

The RNAscope assay produces discrete fluorescent puncta corresponding to individual mRNA transcripts which were counted to quantify levels of *Htr3a* mRNA expression. Each NDNF^+^ and VIP^+^ cell within A1 was imaged at 150x in three channels (e.g. DAPI+*VIP*+*Htr3a*) and photographed using the same exposure times across cases. Both *NDNF* and *VIP* mRNA were highly abundant within cells and used to demarcate cell boundaries. All *Htr3a* puncta contained within these boundaries were manually counted, either by live-counting for low-expressing cells, or from saved images in Fiji for high-expressing cells (>40 puncta per cell). On high-expressing cell images, a standardized rolling ball background subtraction algorithm was performed and brightness/contrast enhanced to ease quantification. A small number of cells were labeled with multiple *Htr3a* puncta without *NDNF* or *VIP* colocalization, possibly corresponding to other categories of 5-HT3R expressing interneurons and were excluded from analysis.

### NDNF Quantification

Cellpose 2.0 human-in-the-loop open-source software was used to automatically segment NDNF puncta using a custom-trained machine learning model (Stringer et al., 2021). Each image was first background subtracted using ImageJ (public domain, Image Processing and Analysis in Java). Cellpose segmented these images to identify individual puncta, then clusters of puncta were manually corrected when grouped inaccurately. A subset of cells expressed high levels of NDNF in which individual puncta were indistinguishable from one another (Figure 4A1). These high-expressing cells were categorized as “saturated” and estimated to have 250 puncta per cell. This number was conservatively chosen to match the highest countable puncta in the unsaturated cases.

### Statistical Analyses

Cell densities were calculated as the number of labeled cells within the ROI divided by the area of the ROI in square millimeters, calculated separately for each cell type within each layer. Cell densities between groups of animals were then compared using the Mann-Whitney U test.

Cellular data for *Htr3a* mRNA were examined using two approaches. First, cells were pooled within each treatment condition and predictions were tested using a generalized linear mixed-effects regression model (GLMM) with a Poisson error distribution. This was chosen because the response variable (number of puncta) was coded as count data which had a Poisson distribution, with most cells expressing low levels of *Htr3a* puncta and a tail of high expressors. NDNF^+^ and VIP^+^ cell populations were analyzed separately, with independent models generated for ontogeny (CTR10 vs. CTR15 vs. CTR30) and for ELS (CTR30 vs. ELS_O_ vs. ELS_R_), for a total of 4 models. We specified condition and layer as fixed effects in each model, with subject and sex as random effects. Models were fit in R (RStudio Team, 2024) using the add-on packages “lme4” (Bates et al., 2015) and “lmerTest” (Kuznetsova et al., 2017). Type III Wald Chi-Square tests were performed on the factors within each model. Post hoc contrast analyses were performed on resulting estimated marginal means using the add-on package “emmeans” (Lenth, 2023). For multiple comparisons, the Benjamini-Hochberg method was used to identify significant p-values at p < 0.05 using a critical value of 0.1. Second, we performed a more conservative analysis of *Htr3a* mRNA data using single animal subject as the unit of analysis. Average *Htr3a* puncta per cell, separated by cell type and layer, were calculated for each animal and subjected to Mann-Whitney U tests to compare developmental trajectories and the effects of ELS.

Cellular data for NDNF puncta quantification were also examined with the two approaches described for *Htr3a* mRNA, using the number of NDNF puncta as the response variable.

Data in text are reported as means ± SEM. In boxplots, purple lines are medians, box edges are 25^th^ and 75^th^ percentiles, and whiskers extend to the most extreme data points excluding outliers.

## RESULTS

### NDNF and VIP Cell Distributions

Consistent with previous reports, we found no colocalization between *NDNF* and *VIP*, which appear as mutually exclusive markers of these cell populations. Across all cases (30 animals), a total of 1,164 NDNF^+^ cells were identified, of which 73.5% resided in L1, accounting for approximately 90% of the labeled cells in this layer. In general, VIP^+^ cells were far more abundant in cortex, with 2,110 cells identified, 2.9% of them residing in L1, 68.3% in L2/3, and 28.8% in L4. Cells in deeper layers were not quantified, as the majority of cortical 5HT3R interneurons are concentrated in L4 and superficial layers (Lee et al., 2010; Kullander and Topolnik, 2021).

### NDNF and VIP Population Densities Decrease During the ACx Critical Period

Both NDNF^+^ and VIP^+^ population densities (# cell bodies/mm^2^) declined precipitously between P10 and P30 under control conditions (Figure 2). A Mann-Whitney U test revealed that NDNF^+^ cell density in L1 was significantly higher in CTR10 (Mean=113.74) compared to CTR15 (M=60.48; *U*=31, *p*=0.04113). A less robust decline in L1 NDNF^+^ cells was seen between CTR15 and CTR30 (M=43.02; *U*=30, *p*=0.06494; Figure 2A). No significant group differences were found between developmental ages in L2/3 or L4 NDNF^+^ cell densities.

**Figure 2.**
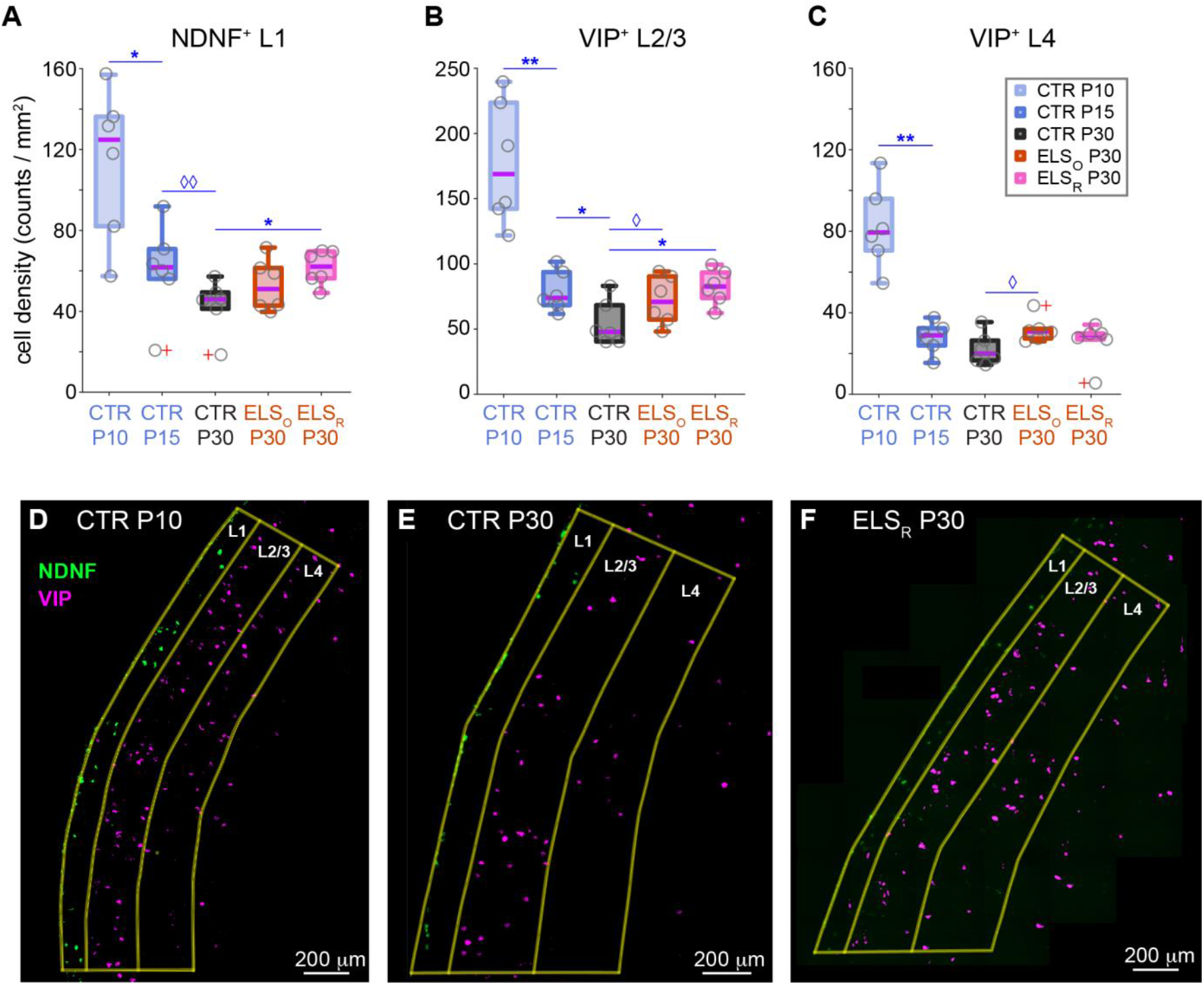
NDNF+ and VIP+ populations drop substantially from P10-P30, but plateau at higher levels under ELS conditions. (A-C) Cell densities within each treatment group; *P* values resulting from pairwise Mann-Whitney U tests (n=6 for all groups). (D-F) Representative A1 maps showing NDNF+ and VIP+ cells for CTR10, CTR30, and ELS_R_. \*\*\*\**p* <0.001, \*\*\**p* <0.002, \*\**p* <0.01, \**p* <0.05, ◊◊*p* <0.08, ◊*p* <0.14. Grey circles in boxplots show individual animal data. Red + symbols inicate outliers.

VIP^+^ cell density in L2/3 was significantly higher in CTR10 than CTR15 (M=78.86; *U*=36, *p*=0.002165; Figure 2B). A significant reduction in L2/3 VIP^+^ density was also observed between CTR15 and CTR30 (M=54.62; *U*=31, *p*=0.04113). In L4, VIP^+^ cell density declines significantly between CTR10 (M=82.19) and CTR15 (M=27.88; *U*=36, *p*=0.002165), but does not differ significantly between CTR15 and CTR30 (M=22.15; *U*=26, *p*=0.2403; Figure 2C). No significant group differences were found for L1 VIP^+^ cell densities.

### ELS Attenuates the Decline of NDNF and VIP Population Densities

Cell population densities for both NDNF^+^ and VIP^+^ were elevated under ELS conditions compared to age-matched controls at P30 (Figure 2A-C). L1 NDNF^+^ cell density was significantly higher in RELS animals (M=61.49) compared to P30 Controls (M=43.02; *U*=32, *p*=0.02597). This effect was not statistically significant when ELS_O_ (M=52.92) were compared to CTR30 (*U*=23, *p*=0.4848). However, the difference was nearly identical to the comparison between ELS_O_ with CTR15 (M=60.48; *U*=23, *p*=0.4848), suggesting that alterations to the ELS_O_ phenotype may be present that escape the sensitivity of this test. L1 NDNF^+^ cell density was similar between CTR15 and ELS_R_ at P30 (*U*=19, *p*=0.9372). NDNF^+^ cell density in L2/3 was also affected by stress, where both ELS_R_ (M=11.77) and ELS_O_ (M=10.43) were significantly higher than CTR30 (M=2.83; *U*=33, *p*=0.02002; *U*=34, *p*=0.01291 for the two comparisons). Within L4, NDNF^+^ density trended higher in ELS_R_ (M=4.26) and ELS_O_ (M=4.97) compared to CTR30 (M=1.66), though this did not reach statistical significance (*U*=30, *p*=0.06367 for each).

L2/3 VIP^+^ cell density was also significantly higher in ELS_R_ (M=82.34) compared to CTR30 (M=54.62), *U*=32, *p*=0.02597. A similar, but less robust trend was seen in ELS_O_ animals (M=71.95), *U*=28, *p*=0.132. Again, neither ELS_R_ nor ELS_O_ differed significantly from CTR15 (*U*=15, *p*=0.6991; *U*=22, *p*=0.5887, respectively). In L4, VIP^+^ cell density did not differ significantly between ELS_R_ (M=25.65) and CTR30, *U*=25, *p*=0.3095, while a difference was trending between ELS_O_ (M=31.78) and CTR30, *U*=30, *p*=0.06494. No significant group differences were found for L1 VIP^+^ cell densities.

### 5HT3a mRNA Expression Increases During Normal Development

For all treatment groups, expression of *Htr3a* mRNA skewed low for NDNF^+^ and VIP^+^ cell populations, with a relatively small number of cells containing high concentrations of labeled puncta (Figure 3C-E). In VIP^+^ cells, *Htr3a* mRNA expression ranged from 0 to 328 puncta, with over 97% possessing 1 or more per cell. NDNF^+^ cells demonstrated a considerably lower expression profile, with *Htr3a* puncta per cell ranging from 0 to 137, and almost 23% containing no *Htr3a*.

**Figure 3.**
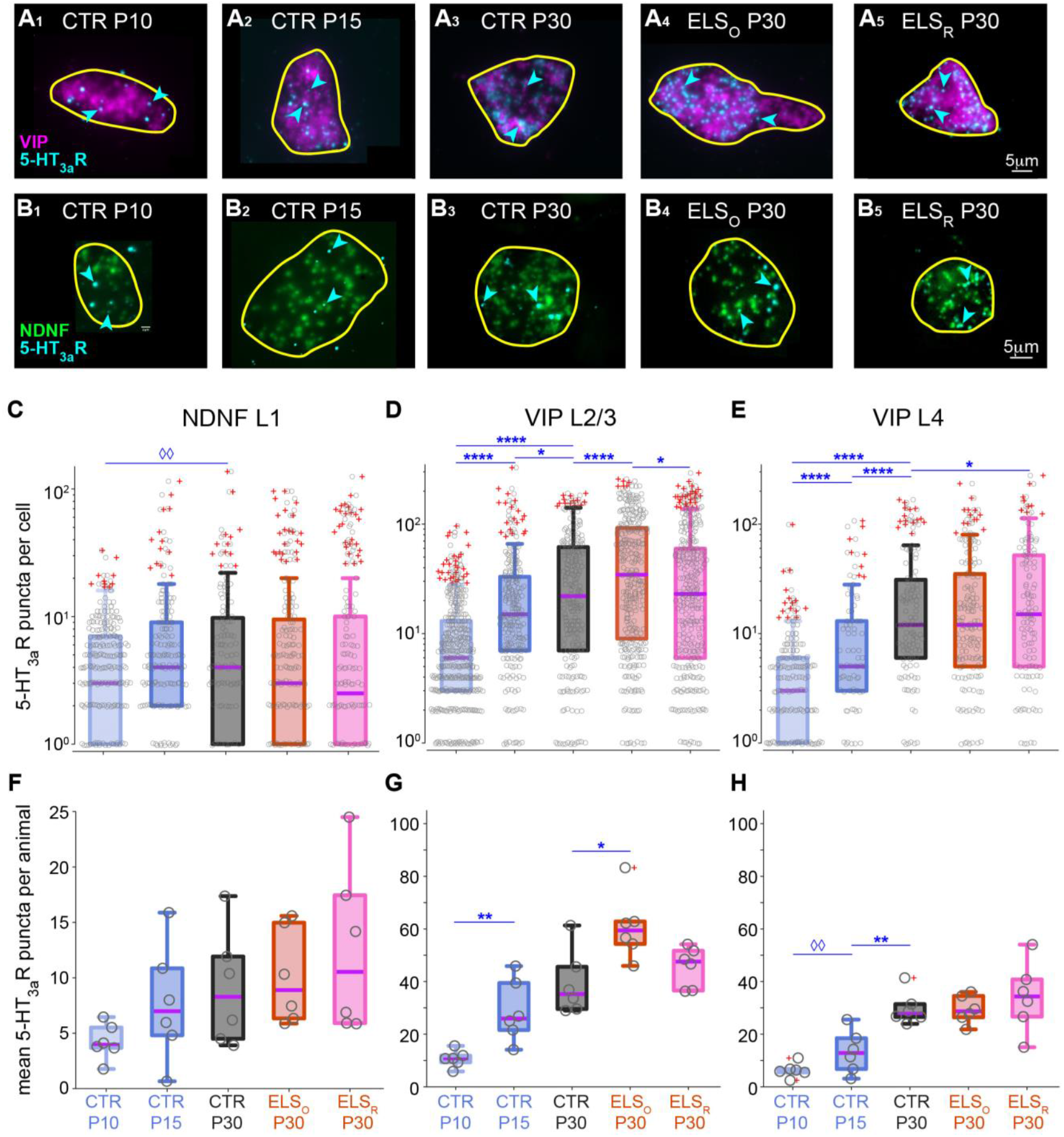
*Htr3a* mRNA increases during the P10-P30 period under control conditions for both cell populations, and is elevated in VIP+ cells by ELS, most prominently in L2/3. (A) L2/3 VIP (*pink*) and *Htr3a* mRNA puncta (*cyan*), in cells representing median expression of *Htr3a* within the highest quartile for each treatment group. (B) L1 NDNF (*green*) and *Htr3a* puncta (*cyan*), in cells representing median expression of *Htr3a* within the highest quartile for each treatment group. (C-E) Log-scale distributions of *Htr3a* puncta per cell within each group. Significant results from the GLMMs are marked above. (F-H) Comparisons of average expression profiles by animal; significant results from Mann Whitney U tests are marked above. \*\*\**p* <0.002, \*\**p* <0.01, \**p* <0.05, ◊◊*p* <0.08. Arrowheads indicate a subset of *Htr3a* mRNA puncta.

We examined the cell populations pooled within each treatment condition through a GLMM approach using four separate models. In our ontogenetic model of VIP^+^ cell development, there was a significant interaction between condition and layer, such that the variance in number of *Htr3a* puncta according to condition depended on layer (Table 2A). Post-hoc tests examining differences between conditions within layers found that L2/3 VIP^+^ cells expressed fewer puncta in CTR10 than CTR15, and fewer puncta in CTR15 than CTR30 (Figure 3D), suggesting a developmental upregulation of the *Htr3a* gene during the P10-P30 period. Similarly, L4 VIP^+^ cells expressed increasing quantities of *Htr3a* mRNA between CTR10, CTR15, and CTR30 (Figure 3E). L1 VIP^+^ cells increased in expression between CTR10 and CTR15, but not significantly between CTR15 and CTR30.

In the ontogenetic model for NDNF^+^ cell development (Table 1B), there was also a significant interaction between condition and layer, though with smaller contrasts, likely related to the lower number of puncta labeled in this population. There was a significant difference in *Htr3a* levels of L1 NDNF^+^ cells between CTR30 and CTR10, but not between CTR15 and other ages (Figure 3C). Number of puncta also increased between P10 and P30 in L2/3 NDNF^+^ cells, but not in L4, where this population is extremely sparse.

**Table 1.**
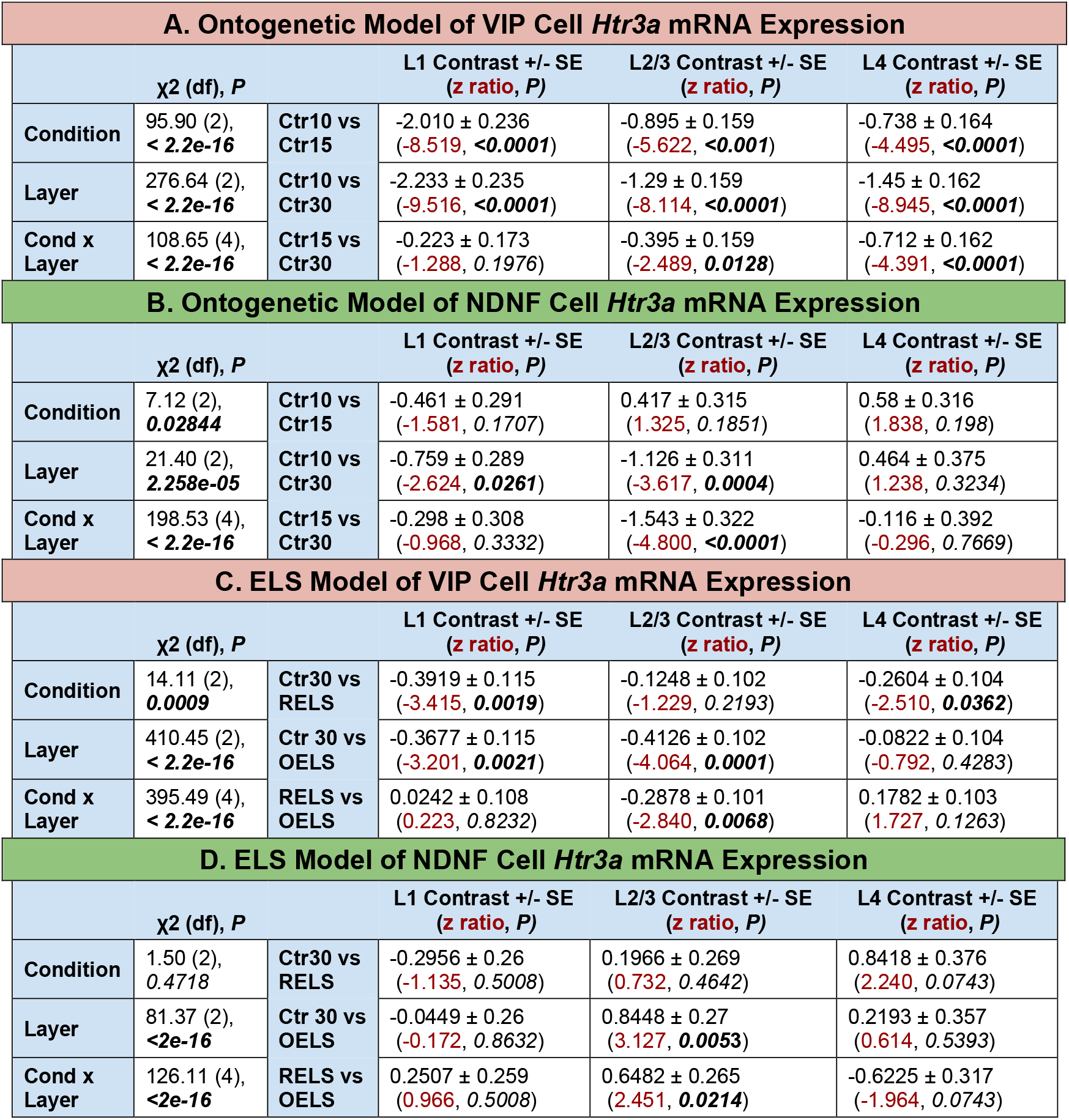
Summary of each of the four generalized linear mixed models (GLMMs) assessing differences in Htr3a mRNA expression. Contrasts were performed on estimated marginal means resulting from the models, with values reported on a log scale. Significant p values with Benjamini-Hochberg adjustment are in **bold**.

To verify the robustness of these results, we calculated the average puncta per cell for VIP^+^ and NDNF^+^ populations within layers for each animal then subjected the averages to Mann-Whitney U tests. Under control conditions, we found that average *Htr3a* puncta per cell increases during the course of normal development in VIP^+^ cells (Figure 3G,H). In L2/3 VIP^+^ cells, mRNA expression is significantly higher in CTR15 (M=28.82) compared to CTR10 (M=11.14) (*U*=35, *p*=0.0043). This trend continued between CTR15 and CTR30 (M=39.89), but did not reach significance (*U*=27, *p*=0.1797). Similar increases were seen in L4 VIP^+^ cells (Figure 3H). Average puncta per cell trended upward between CTR10 (M=6.56) and CTR15 (M=13.48) (*U*=29, *p*=0.09307), then increased significantly between CTR15 and CTR30 (M=28.80) (*U*=34, *p*=0.0087). No significant group differences were found between control groups for L1 VIP^+^ cells.

For NDNF^+^ cells, though there is a trend toward increasing average expression, we did not find any significant group differences (Figure 3F).

### ELS Increases 5HT3a mRNA Expression in VIP^+^ Cells in a Layer-Specific Manner

Similar to the ontogenetic models, our ELS model of VIP^+^ cells revealed a significant interaction between condition and layer, with contrasts showing distinct layer-specific effects of each stress group (Table 1C). In the L2/3 VIP^+^ population, *Htr3a* was significantly higher in ELS_O_ compared to CTR30, whereas there was no significant contrast between ELS_R_ and CTR30 (Figure 3D). Conversely, among L4 VIP^+^ cells, mRNA expression in ELS_R_ was higher than CTR30, with no significant difference between ELS_O_ and CTR30 (Figure 3E). Both ELS_O_ and ELS_R_ contained significantly more *Htr3a* puncta in L1 compared to CTR30, though these cells are few.

The ELS model of NDNF^+^ cells identified a significant interaction between layer and condition but did not identify condition on its own as a significant predictor of *Htr3a* puncta levels, diverging from the other models (Table 1D). No differences were found in L1 NDNF^+^ cells. Indeed, the only significant contrast found was between ELS_O_ and ELS_R_ L2/3 NDNF^+^ cells, sparse populations containing only 64 and 51 cells, respectively.

Comparing averages between animals, we found that *Htr3a* mRNA expression in L2/3 VIP^+^ cells is greater in ELS_O_ (60.86, n=6) compared to CTR30 (M=39.89) (*U*=33, *p*=0.0152), while ELS_R_ (M=45.65) did not differ significantly from controls (*U*=26, *p*=0.2403) (Figure 3G). Mann-Whitney U tests did not reveal any significant differences between ELS_R_ or ELS_O_ with controls in L1 or L4 VIP^+^ cells. No significant group differences were found between stress groups and controls for NDNF^+^ cells in any layers.

### NDNF mRNA Expression Decreases During Normal Development

To evaluate whether development or stress affected levels of NDNF mRNA within NDNF^+^ cells, we quantified *NDNF* in all NDNF cells across layers 1 through 4. Similar to our approach with *Htr3a* data, we examined *NDNF* expression pooled within each treatment condition using two separate GLM models for normal development and for evaluation of ELS effects. In our ontogenetic model of NDNF^+^ cell development, there was a significant interaction between condition and layer, such that the variance in number of *NDNF* puncta according to condition depended on layer (Table 2A). Post-hoc tests examining differences between conditions within layers found that L1 cells expressed significantly more puncta in Ctr10 than Ctr30 (Figure 4B), suggesting a developmental downregulation of *NDNF* expression during this window. This trend was also noted between Ctr10 vs Ctr 15 animals, as well as Ctr15 vs Ctr30 animals, but did not reach significance in these individual contrast analyses. No developmental differences were seen in L2/3 NDNF^+^ cells. L4 contrasts yielded mixed results: a non-significant decline in expression was observed between Ctr10 to Ctr15, accompanied by an unexpected significant increase in expression between Ctr15 to Ctr30. As there were relatively few NDNF^+^ cells in L4 compared to L1, and no previous description of L4 NDNF^+^ neurons in the existing literature to our awareness, we interpret these findings cautiously.

**Figure 4.**
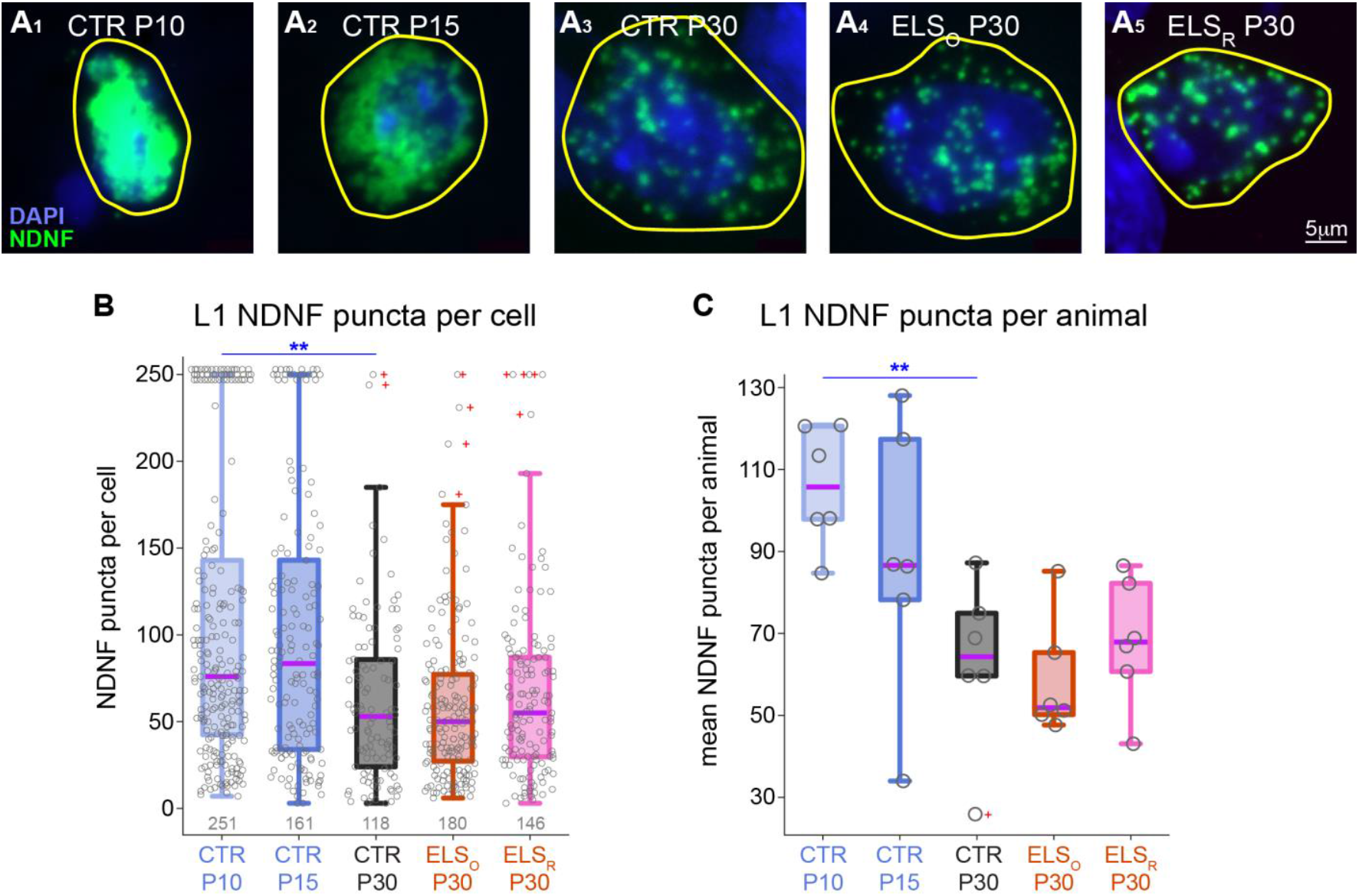
Layer 1 NDNF neurons show a developmental decrease of *NDNF* mRNA. (A) L1 *NDNF* mRNA puncta (*green*), in cells representing median expression of *NDNF* within the highest quartile for each treatment group. (B) Distributions of L1 *NDNF* puncta per cell within each group. Significant results from the GLMMs are marked above. Grey numbers indicate number of cells per treatment group. Grey circles in boxplots show individual cell data. (C) Within L1 NDNF cells, comparisons of average *NDNF* expression profiles by animal; significant results from Mann Whitney U tests are marked above. Grey circles in boxplots show individual animal data. \*\*\*\**p* <0.001, \*\*\**p* <0.002, \*\**p* <0.01, \**p* <0.05, ◊◊*p* <0.08, ◊*p* <0.14.

In line with our treatment of *Htr3a* data, we calculated the mean *NDNF* puncta per cell within L1 for each animal and subjected these averages to Mann-Whitney U tests. Under control conditions, we again observed a decline in *NDNF* expression during normal development (Figure 4C). mRNA expression was significantly greater in Ctr10 (M=105.9) compared to Ctr30 (M=62.71) (*U*=35, *p*=0.004329). There was a consistent downward trend from Ctr10 to Ctr15, and between Ctr15 to Ctr30, though these individual comparisons did not reach significance, consistent with the outcome of our GLMM. No significant differences were found within L2/3 or L4 under normal development.

### NDNF mRNA Expression is Mostly Unaffected by ELS

The ELS model of NDNF^+^ cells identified a significant interaction between layer and condition but did not identify condition on its own as a significant predictor of NDNF puncta levels (Table 2B). No significant contrasts were found between conditions for L1 (Figure 4B) or L2/3 NDNF^+^ cells. In L4, NDNF expression was significantly greater in RELS animals compared to P30 controls, whereas expression was significantly lower in OELS animals compared to P30 controls. The largest difference was observed between the stress conditions themselves, with significantly greater L4 expression levels in RELS compared to OELS.

Comparing averages between animals, no significant effects of stress emerged in either L1 (Figure 4C) or L2/3. In L4, similar to our GLMM analyses, RELS animals (M=92.5) showed greater expression compared to Ctr30 (M=40.5) (*U*=15, *p*=0.036) and to OELS (M=36.3) (*U*=29, *p*=0.0087). There was no difference found between OELS vs. Ctr30 subjects. Again, because of the low number of cells sampled within this population, and with some animals lacking L4 NDNF data altogether, these findings should be interpreted with caution as they may be spurious due to insufficient sampling.

**Table 1a.**
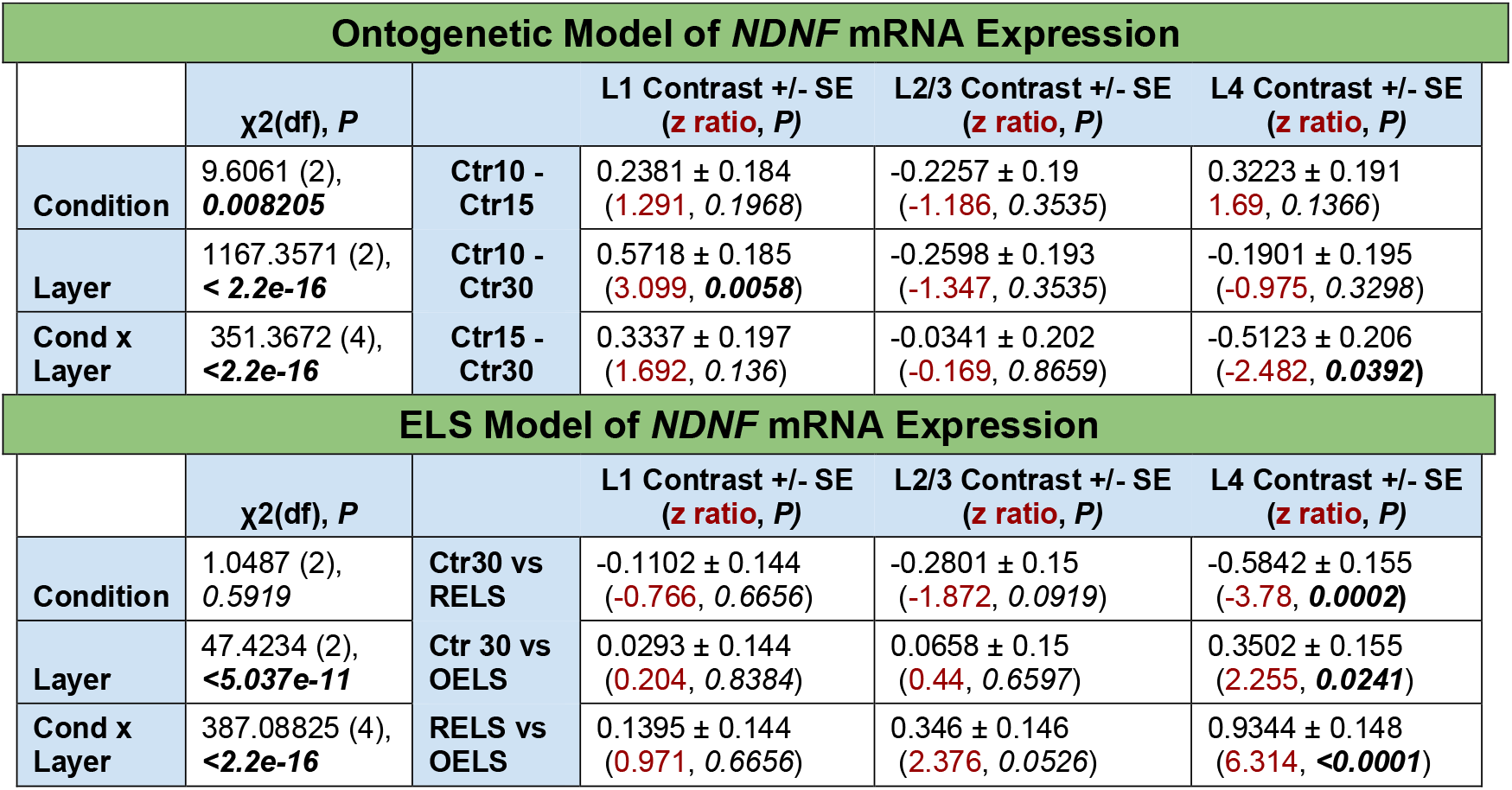
Summary of the two GLMMs assessing differences in NFNF mRNA expression. Contrasts were performed on estimated marginal means resulting from the models, with values reported on a log scale. Significant p values with Benjamini-Hochberg adjustment are in **bold**.

## DISCUSSION

5-HT3R expressing inhibitory interneurons, particularly L1 NDNF^+^ and L2/3 VIP^+^ cells, have gained increased attention in recent years as important regulators of cortical circuitry. These superficial neurons influence developmental plasticity and integrate top-down and bottom-up information to gate response properties of other cortical neurons (Takesian et al., 2018; Pardi et al., 2023). The gene encoding the 3A protein subunit, *Htr3a*, present in all functional 5-HT3Rs (Boyd et al., 2002), has been identified as a genetic locus that is susceptible to ELS and may be linked to clinical outcomes in neuropsychiatric conditions (Rajkumar and Mahesh, 2010; Gupta et al., 2016; Perroud et al., 2016). By examining the *in situ* cell-specific expression of *Htr3a* across the ACx critical period, our data provide context for how ELS may interact with this gene to produce functional aberrations in the circuits involving NDNF^+^ and VIP^+^ interneurons.

Developmentally we found that in ACx, VIP^+^ and NDNF^+^ cell densities decreased dramatically between P10 and P30 in control animals, with much of the reduction in cell density occurring by P15, midway through the ACx critical period and following ear opening at ∼P11 (Figure 2). For both cell types, CTR10 animals showed substantial variation in cell density that largely collapsed by P15, indicating that at this time point we sampled a rapid shift in the development of the cell populations. The timing of this decrease following the increased sensory input of ear opening is suggestive of an activity-dependent process and may reflect apoptosis. Developmental apoptosis has not been described in auditory cortex, but in the auditory brainstem, hearing onset precedes a period of cell death (Tierney and Moore, 1997), and apoptosis across cortical regions is regulated based on activity levels (Blanquie et al., 2017). Previous work has established that interneurons derived from the caudal ganglionic eminence (CGE), which constitute the bulk of 5-HT3R cells, undergo programmed cell death, and that this occurs in an activity-dependent manner that relies on calcineurin signaling during specific periods in postnatal development (De Marco Garcia et al., 2011; Priya et al., 2018). VIP interneurons have been singled out as an exception to this rule, with some experiments suggesting that they are exempt from this activity-dependent effect. However, more recent work discovered that different mechanisms govern the survival of distinct 5-HT3R subpopulations, with VIP interneuron fate specifically tied to serotonergic input during active periods of programmed cell death (Wong et al., 2022). The ontogeny of serotonin metabolism in ACx is consistent with this result, increasing between P10 and P15 in rats (Basura et al., 2008), and the 5-HT3Rs present in these cells are excitatory. Thus the sharp decline in densities of both of these inhibitory cell types may reflect cell death that is cued by the increased activity accompanying ear opening.

Our data show that ELS attenuates the developmental decline in both NDNF^+^ and VIP^+^ cell density in a layer-specific manner (Figure 2). In ELS animals, we found increased cell density in L2/3 VIP^+^ cells at P30 compared to controls. In ELS_O_ animals, these cells also expressed higher levels of *Htr3a* (Figure 3), raising the possibility that ELS may impact VIP^+^ cell survival through upregulation of the 5-HT3A gene, a possibility consistent with a known protective role of 5-HT in cell survival (Vitalis and Parnavelas, 2003; Barzegar-Fallah et al., 2023), though that action may be context dependent (Ju Yeon and Yeon Hee, 2005; Aminzadeh, 2017). Similar effects were seen in prefrontal cortex, where ELS in rats increased the density of VIP cells (Pascual et al., 2006). Further work is needed to confirm the specific contributions of 5-HT3A, given the milieu of changes brought about by ELS on the serotonin system (Lages et al., 2021; Lipsky et al., 2022). Similarly, L1 NDNF^+^ cell density was also significantly increased by stress, with P30 ELS_R_ animals maintaining cell populations on par with P15 controls. These cells, however, did not show a marked increase in *Htr3a* expression. It is possible that other changes, such as glutamatergic inputs known to influence interneuron cell survival, may instead influence NDNF cell fate under these conditions (Wong et al., 2022).

The results of our GLMM analyses revealed that *Htr3a* mRNA was developmentally upregulated in VIP^+^ cells in all layers that were examined (L1, L2/3, and L4), whereas levels remained relatively stable in NDNF^+^ cells (Figure 3; Table 1). Note that *Htr3a* expression was ∼four-fold higher in VIP^+^ than NDNF^+^ cells, consistent with a stronger depolarizing effect of serotonin on VIP^+^ cells mediated at least in part by 5-HT3Rs (Poorthuis et al., 2018; Pronneke et al., 2020; Wong et al., 2022). Stress induction increased *Htr3a* expression beyond control levels in VIP^+^ cells, with ELS_O_ animals showing substantial increases in L2/3, and ELS_R_ animals showing increased levels in L4. While it is unclear why the two stress induction methods used here would not cause equivalent changes, it is well established that ELS effects can vary based on many parameters including type of stressor, animal age at induction, duration and unpredictability of stress, and individual genetic susceptibility (Lehmann and Feldon, 2000; Rincon-Cortes and Sullivan, 2014). Neither form of ELS had a substantial impact on *Htr3a* expression in NDNF^+^ cells. These findings were largely recapitulated when using single animal as the unit of analysis. Though the extent to which elevated *Htr3a* mRNA translates to increased protein production is unknown, differentially expressed genes generally show greater correlations between mRNA and protein quantity compared to genes that maintain stable expression profiles (Koussounadis et al., 2015). Increased *Htr3a* is consistent with several studies in which developmental stress altered 5-HT3R expression in cortical regions, though these approaches did not evaluate cell-type specificity (Perroud et al., 2016; Moradi-Kor et al., 2020; Zolfaghari et al., 2021).

Increased expression of functional excitatory 5-HT3Rs over development and by early-life stress, combined with increased NDNF^+^ and VIP^+^ numbers, could alter both cortical plasticity and auditory processing and contribute to the temporal processing deficits that follow ELS. Both VIP^+^ and NDNF^+^ interneurons integrate inputs from primary auditory thalamus and neuromodulatory regions (Kuchibhotla et al., 2017; Poorthuis et al., 2018; Vattino et al., 2025), responding with high selectivity to relevant or novel features of complex auditory stimuli (Zhu et al., 2015; Cohen-Kashi Malina et al., 2021; Rikhye et al., 2021; Yarden et al., 2022; Jamali et al., 2024; Sweeney et al., 2025), particularly in specific behavioral contexts (Ding et al., 2025; Sweeney et al., 2025). The involvement of these neurons in context-dependent responses to temporally-varying behaviorally relevant sounds may be disrupted by ELS-induced circuit changes. Inhibitory fibers from VIP^+^ cells target a combination of intracolumnar cell bodies of PV and somatostatin (SST) interneurons (Figure 1a), forming disinhibitory circuits that increase pyramidal cell activity (David et al., 2007; Hioki et al., 2013; Pi et al., 2013; Zhang et al., 2022). Increased expression of the excitatory 5-HT3R along with increased numbers of VIP^+^ cells could thus enhance pyramidal excitability, though this may be counteracted by circuit effects involving SST interneurons (Batista-Brito et al., 2017; Ferguson et al., 2023). NDNF^+^ cells, though fewer in number than VIP^+^, are regularly spaced throughout L1 and exert a powerful influence over cortical dynamics, integrating top-down signals with bottom-up sensory input to gain-modulate cortical columns (Cohen-Kashi Malina et al., 2021; Vattino et al., 2025). They receive inputs from widespread sources whose fibers course through L1, and through volume transmission coordinate a mixture of postsynaptic targets that include PV cells, VIP^+^ cells, and pyramidal cell apical dendrites to affect state-dependent sensory processing (Letzkus et al., 2011; Overstreet-Wadiche and McBain, 2015; Hartung et al., 2024; Sweeney et al., 2025). Both cell types have been implicated in plasticity (Takesian et al., 2018; Melzer et al., 2021; Williams et al., 2025). Developmentally, shifts in NDNF^+^ and VIP^+^ activity during the ACx critical period may dramatically alter circuit dynamics. Indeed, developmental perturbation of NDNF^+^ or VIP^+^ activity produced perceptual deficits along with altered cortical map organization and response properties of pyramidal cells, including sensory-evoked firing, phase-locking, and response selectivity (Batista-Brito et al., 2017; Che et al., 2018; Takesian et al., 2018), indicating a role for these cells in developmental plasticity. Thus alterations to NDNF^+^ and VIP^+^ dynamics by stress, via a combined action of increased sensitivity to serotonin and increased cell survival, could lead to imbalanced excitation within affected cortical columns, impairing perception and learning.

Because both VIP^+^ and NDNF^+^ cells target PV interneurons (Pi et al., 2013; Hartung et al., 2024), PV cells may be affected by changes to 5-HT3R interneurons. PV neurons contribute to accurate temporal processing and drive critical period plasticity (Krause et al., 2019; Reh et al., 2020; Nocon et al., 2023), both of which are shifted by ELS (Pattwell and Bath, 2017; Ye et al., 2023; An et al., 2025; Mazi et al., 2025; Rosen and Huyck, 2025). ELS is known to affect PV cells, decreasing PV expression and reducing cell density in ACx and other regions (Goodwill et al., 2018; Sunthimer et al., 2023; An et al., 2025; Mazi et al., 2025). Further, cell survival in interneurons derived from the medial ganglionic eminence (MGE), including PV cells, is affected by levels of cell activity (Denaxa et al., 2018). Thus PV survival may be reduced by increased inhibitory tone from elevated NDNF^+^ and VIP^+^ cell populations containing more excitatory 5-HT3Rs, leading to disruption of inhibitory-excitatory dynamics. It is worth noting that because the expression of PV is activity dependent, reduced PV cell density indicated by antibody binding assays may in part reflect lowered activity in PV neurons. During development, reduced PV associates with delayed critical period plasticity (Koh and Sng, 2016; Reh et al., 2020), and auditory temporal processing deficits can be rescued by PV activation (Masri et al., 2021; Masri et al., 2024). PV activity enables accurate responses to temporally-varying stimuli (Nocon et al., 2023); for example PV cells respond more strongly to short gaps than pyramidal cells (Keller et al., 2018). Thus PV reduction by ELS could contribute to the ELS-induced temporal processing deficits that have been recently described in gerbils, rats, and mice (Ye et al., 2023; An et al., 2025; Mazi et al., 2025).

Finally, we examined expression of *NDNF* mRNA within NDNF^+^ cells to reveal a developmental decline in *NDNF* that was not affected by ELS. NDNF is a secretory protein that promotes cell survival, migration and neurite outgrowth, while protecting against excitotoxity and downregulating apoptosis-related pathways (Kuang et al., 2010; Cheng et al., 2023a; Liu et al., 2024). While it is not established whether NDNF^+^ L1 neurons have a secretory role, their propensity for volume transmission suggests that these cells could provide broad trophic support to their targets (Olah et al., 2009). The large drop in *NDNF* mRNA from P10 to P30 coincides with ACx critical period plasticity and increased activity following ear opening, a period when cortical circuitry is refined based on activity levels and trophic support from neural growth factors (Catapano et al., 2001). At P10 we identified many cells containing a very high density of NDNF puncta (Figure 4B); this declined over development and rarely occurred by P30. Although this study did not evaluate the role of NDNF as a trophic factor, the gradual decline of NDNF coincides with reduced NDNF^+^ and VIP^+^ cell density. However, NDNF mRNA levels remained stable with ELS, despite increased numbers of NDNF^+^ and VIP^+^ cells. Thus NDNF may not contribute to the increased NDNF^+^ and VIP^+^ cell densities seen with ELS. It is possible that other trophic factors and/or increased activity levels in ELS cells, predicted by increased 5HT3aRs, contributes to the higher number of cells following ELS.

The present study identified changes to 5-HT3R cell populations as a function of development and early-life stress, including an increase in cell density and *Htr3a* expression following ELS. The functional implications of these differential expression profiles will need to be evaluated by assessing the responsiveness of these subpopulations to serotonergic input and measuring the behavior of their synaptic targets. Future studies are needed to reveal the full range of functional sequelae resulting from these effects and shed further light on potential interventions to mitigate such maladaptive alterations during development.

## Acknowledgements

This work was supported by NIDCD R01 DC013314 to MJR, NIDCD R01 supplement DC013314-09S1 to JTM, and NIDCD R01 DC017708 to JGM. The content is solely the responsibility of the authors and does not necessarily represent the official views of the National Institutes of Health. The authors would like to thank Dr. Jesse W. Young for help with statistical modeling, and Dr. Jeffrey Wenstrup for comments on an earlier version of the manuscript.

## Notes

### Competing Interest Statement

The authors have declared no competing interest.

### Summary of Updates

This revision fixes a document conversion error that had caused a table and figure to obscure one another.

